# A new model of chronic *Mycobacterium abscessus* lung infection in immunocompetent mice

**DOI:** 10.1101/2020.07.30.228247

**Authors:** Camilla Riva, Enrico Tortoli, Federica Cugnata, Francesca Sanvito, Antonio Esposito, Marco Rossi, Anna Colarieti, Tamara Canu, Cristina Cigana, Alessandra Bragonzi, Nicola Ivan Loré, Paolo Miotto, Daniela Maria Cirillo

## Abstract

Pulmonary infections caused by *Mycobacterium abscessus* (MA) have increased over recent decades affecting individuals with underlying pathologies such as chronic obstructive pulmonary disease, bronchiectasis and, especially, cystic fibrosis. The lack of a representative and standardized model of chronic infection in mice has limited steps forward in the field of MA pulmonary infection. To overcome this challenge we refined the method of agar beads to establish MA chronic infection in immunocompetent mice. We evaluated bacterial count, lung pathology and markers of inflammation and we performed longitudinal studies with magnetic resonance imaging (MRI) up to three months after MA infection. In this model, MA was able to establish a persistent lung infection up to two months and with minimal systemic spread. Lung histopathological analysis revealed granulomatous inflammation around bronchi characterized by the presence of neutrophils, lymphocytes and aggregates of vacuolated foamy cells, mimicking the damage observed in humans. Furthermore, MA lung lesions were successfully monitored for the first time by MRI. The availability of this murine model and the introduction of the successfully longitudinal monitoring of the murine lung lesions with MRI pave the way for further investigations on the impact of MA pathogenesis and the efficacy of novel treatments.

## 1. Introduction

Nontuberculous mycobacteria (NTMs) are environmental organisms ubiquitous in water and soil. The large majority of NTMs are not pathogenic to humans, causing disease only in the presence of predisposing host conditions [1]. The prevalence of pulmonary NTM infections has increased over recent decades [2] affecting individuals with preexisting lung inflammation and compromised ability to clear the infection such as in chronic obstructive pulmonary disease (COPD), bronchiectasis and, especially, cystic fibrosis (CF) disease [3]. The prevalent NTMs infecting CF patients are members of *Mycobacterium avium* complex and *Mycobacterium abscessus* (MA) that includes three subspecies (subsp.), *MA* subsp. *abscessus, MA* subsp. *bolletii* and *MA* subsp. *massiliense* [4]. MA is a rapidly growing NTM responsible for chronic pulmonary infections associated with diversified clinical presentations ranging from asymptomatic colonization to a significant decline of lung function associated with morbidity and mortality, and with poor treatment outcome. Despite prolonged therapy with multiple antibiotics used in different combinations, MA infections are extremely difficult to eradicate due to the multidrug resistance of the mycobacterium [5, 6], that restricts the number of molecules available for the treatment. Among the multiple cellular and animal models proposed for studying the pathology of MA infection, very few could be used to evaluate new drugs efficacy [7]. Murine and human primary macrophages or other cell lines were used to dissect the early MA invasion of phagocytic cells. Amoebae were useful as host models to study and identify the expression of several determinants that contribute to MA virulence [7]. Zebrafish (*Danio rerio*) embryos were exploited during the last two decades to study the intricate interactions between MA and the host immune system. Immunocompetent mouse models of infection following aerosol [8], intratracheal [9], or intravenous [10] bacteria inoculation, were characterized by transient colonization with a rapid clearance of MA in the first weeks post challenge. Alternative models based on immunocompromised murine models were developed (e.g. GM-CSF -/-, GKO or SCID mice) [11-13] which have resulted in slower resolution of MA lung infection than immunocompetent mice. Despite these advancements, the lack of a standardized mouse model of persistent chronic infection in immunocompetent mice that reflects the human MA lung pathology, is the major impediment to studying MA pulmonary infection and treatment strategies [14]. In this study, we used *MA* subsp. *abscessus, MA* subsp. *bolletii* and *MA* subsp. *massiliense* to establish a long-term chronic pulmonary infection in immunocompetent mice. To infect mice we adapted to MA a method based on intratracheal inoculation of bacteria embedding agar beads, previously used with other pathogens (*Pseudomonas aeruginosa, Staphylococcus aureus* and *Burkholderia cenocepacia*) [15-17]. We monitored and analyzed the host response (bacterial burden, pulmonary lesions and inflammatory cytokines) for up to 90 days from infection. Furthermore, we explored the possibility to monitor longitudinally the MA lung challenge within the same animal by using the magnetic resonance imaging (MRI) technique for the first time.

## 2. Results

### 2.1 Chronic lung persistence of MA in C57BL/6NCrl mice

MA subsp. *abscessus* embedded in the agar beads was intratracheally inoculated in immunocompetent C57BL/6NCrl mice to generate an effective long-term chronic lung infection. Different groups of mice were sacrificed at different time points (7, 14, 28, 45, 65, 90 days). Results showed that up to 65 days after infection the overall rate of MA subsp. *abscessus* chronicity was 97.56% (**Figure 1A**). The colony forming units (CFUs) were not significantly different among the time points 7, 14, 28, 45 and 65 (Kruskal-Wallis test p-value=0.0752) with a stable bacterial load (median: ∼10^6^ CFUs) (**Figure 1B**). After this time point, some mice started clearing the bacteria and the lung chronicity rate significantly decreased to 40% (**Figure 1A, Table S1**). MA *bolletii* (**Figure S2A** and **B, Table S3**) and MA *massiliense* (**Figure S3A** and **B, Table S5**) were chronically infected for up to 45 days, but after 7 days of infection the number of CFUs started to decrease with statistical significance. Of note, lung infection of the 3 subspecies did not lead to a systemic infection as indicated by the number of CFU observed in the spleen (**Figure S1** and **Table S2**; **Figure S2C** and **Table S4**; **Figure S3C** and **Table S6**). The body weight (**Figure S4A, Table S7**) and the Kaplan-Meier survival curves (**Figure S4B, Table S8**) of the 3 subspecies did not show differences in comparison with control mice, injected with empty beads.

**Figure 1.**
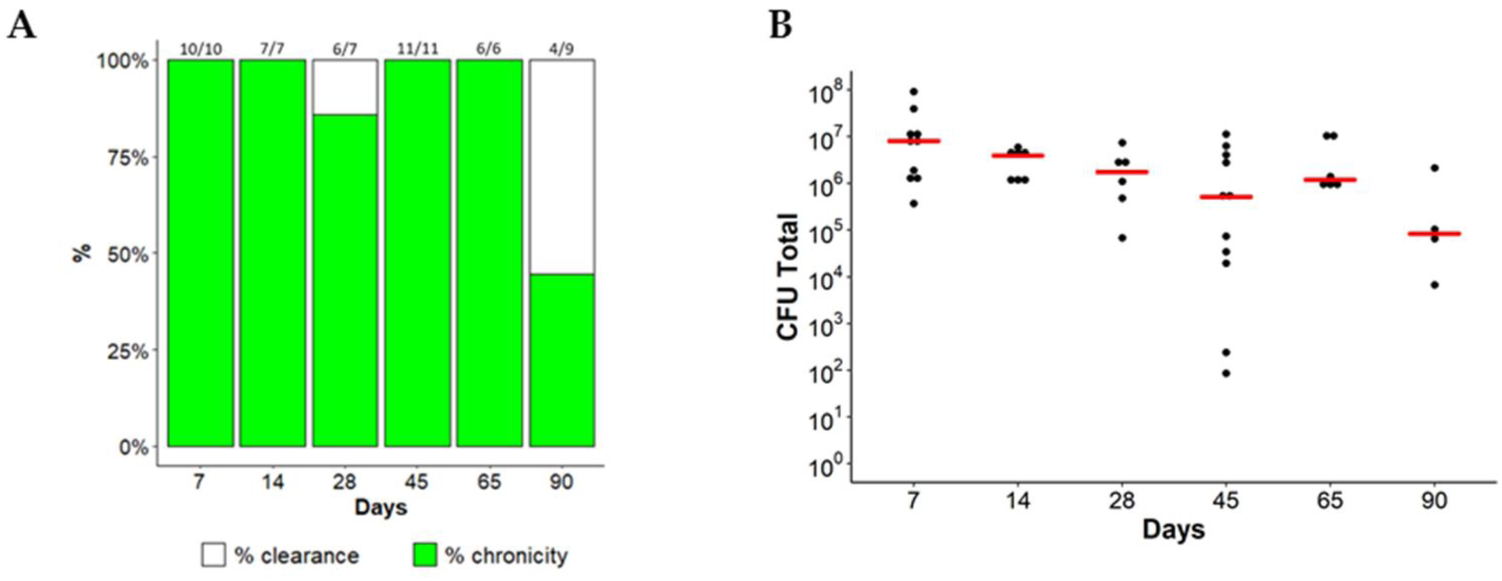
Time course of MA subsp. *abscessus* chronic lung infection. C57BL/6NCrl mice were intratracheally infected with 1×10^5^ of MA *abscessus* reference strain (ATCC 19977) embedded in agar-beads. After 7, 14, 28, 65, 90 days from infection mice were sacrificed. (**A**) Rate of chronicity (number of mice = 6-11). The data were pooled from two independent experiments (**Table S1**). (**B**) Mice were considered infected with CFUs >0 in total lung. Dots represent CFUs in individual mice. The red line represents the median values. Kruskal-Wallis test was performed.

### 2.1 Lung lesions and inflammatory response during MA chronic infection in C57BL/6NCrl mice

Histopathological characterization of lungs infected by MA subsp. *abscesuss* was performed at different time points for 3 months. Morphological analysis after 7 days of infection revealed aggregates of inflammatory cell infiltrate in the parenchyma, characterized by IBA-1 positive-histiocytes and foamy cells, surrounded by few CD3 and few B220 positive lymphocytes (**Figure 2A**). After 45 days of infection the inflammatory cell infiltrate was organized as granulomas mainly localized in the parenchyma around bronchi and characterized by a central core of IBA-1 positive-histiocytes intermingled with granulocytes, surrounded by, CD3 and B220 positive-lymphocytes (**Figure 2A**). Microscopic analysis after 90 days showed a residual peribronchial cuffing of CD3, B220, and IBA-1-positive cells (**Figure 2A**). An example of MA subsp. *abscessus* lung localization after 45 days of infection is reported in supplementary data (**Figure S5**, Ziehl Neelsen staining). The quantification of the total lung lesions (inflammatory cell infiltrate, granulomas and bleeding area) induced by the MA subsp. *abscessus* was statistically higher than in control mice, during the course of the infection (**Figure 2B, Table S9**). Similar results were obtained with MA subsp. *bolletii* and MA subsp. *massiliense* (**Figure S6A and B, Table S10** and **Table S11**). TNF-α (**Figure 3A**) and IFN-γ (**Figure 3B**) showed a significant peak expression at an early time point of MA subsp *abscessus* infection compared to control mice. These results were confirmed for MA subsp. *bolletii* (**Figure S7**) and MA subsp. *massiliense* (**Figure S8**).

**Figure 2.**
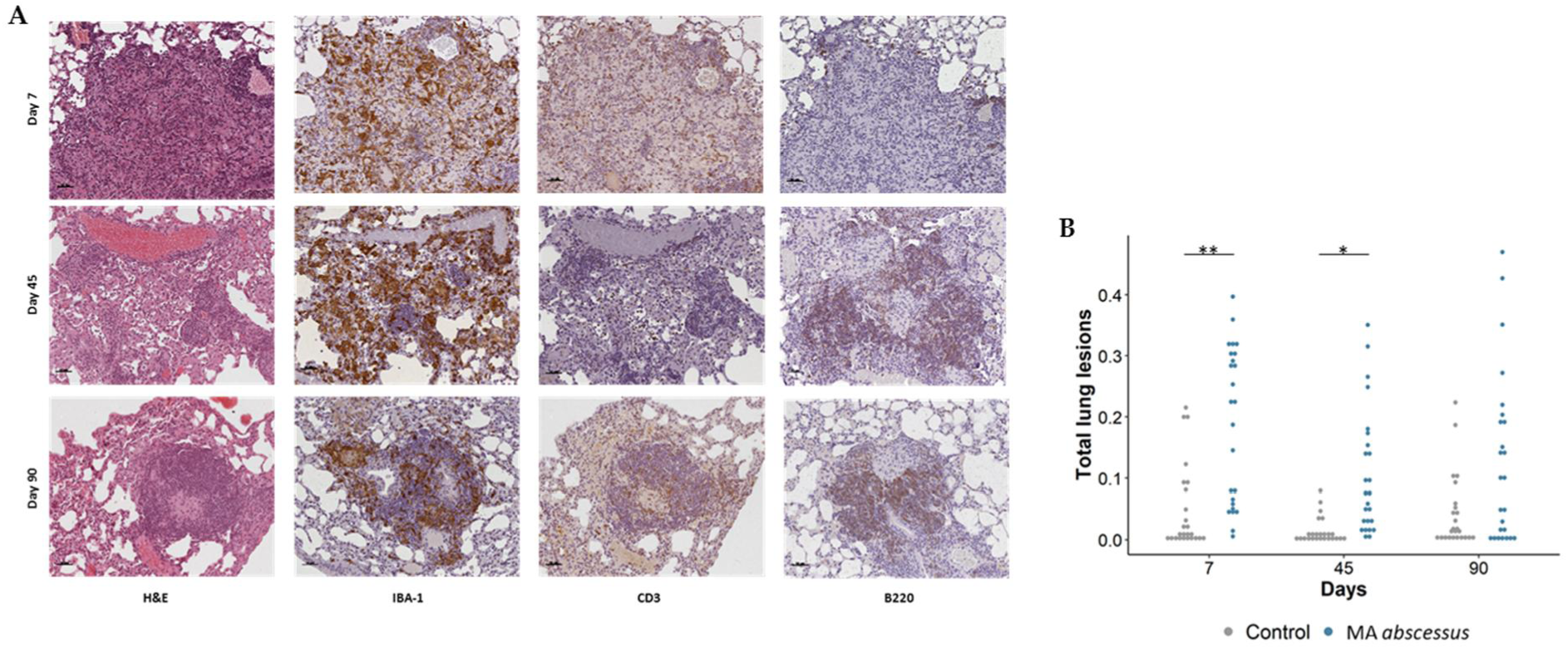
Lung lesions after 7, 45 and 90 days of chronic infection with MA subsp. *abscessus*. (**A**) The panel shows H&E, anti-IBA 1, anti-CD3 and anti-B220 (20X, AxioCam HRc Zeiss) stained sections of lungs after 7, 45 and 90 days from infection; scale bar: 50 µm. (**B**) Dots represents the lobes analyzed by Image J program: 5 lobe images of 5 mice for each time point (25 dots for each time point). Total lung lesions were estimated in mice infected or not by MA subsp. *abscessus*. Linear mixed model followed by post hoc analysis was performed (**Table S9**). Significance was calculated for each time point comparing MA subsp. *abscessus* and control. *p<0.05, **p<0.01, ***p<0.001 ****p<0.0001.

**Figure 3.**
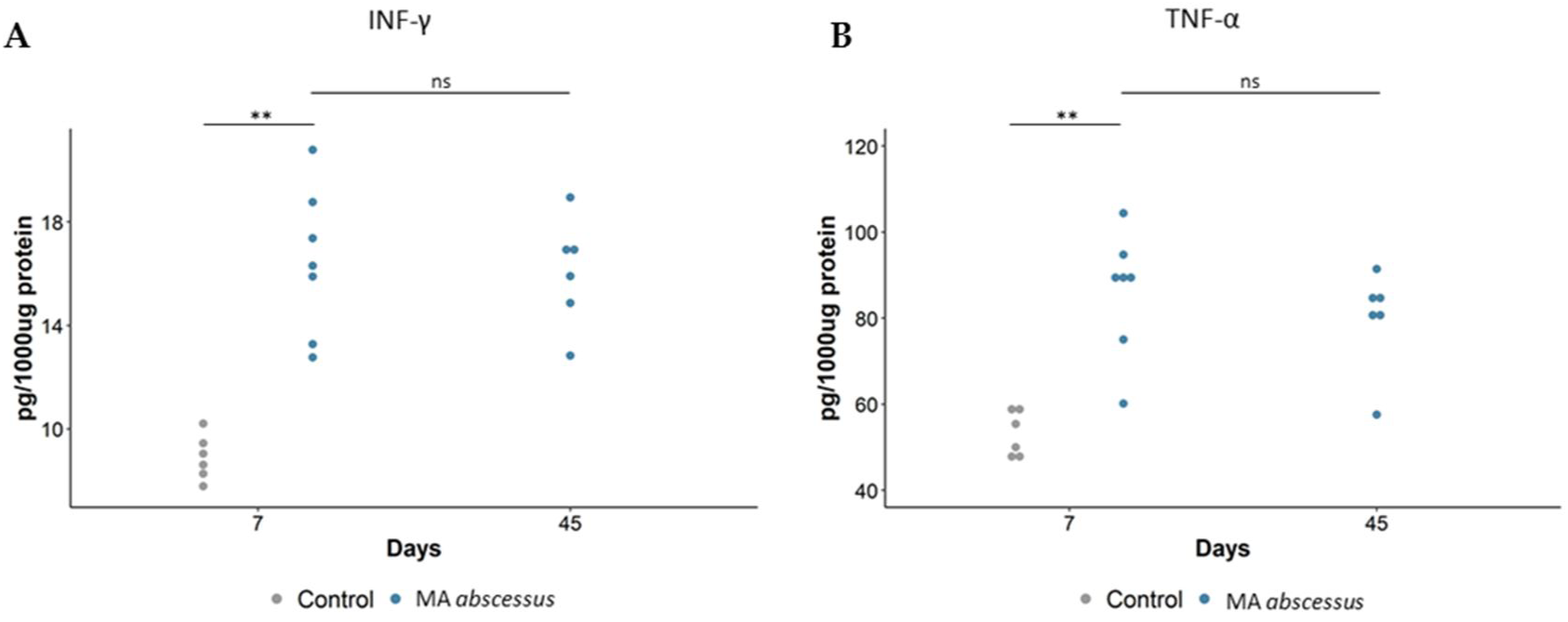
Cytokines/chemokines recruitment during MA subsp. *abscessus* chronic lung infection. (**A**) IFN-γ, (**B**) TNF-α, measured by Mouse Milliplex, were quantified in total the lung of mice. At 7 and 45 days, dots represent cells in individual mice selected from the group of infected mice with MA subsp. *abscessus*. The data were pooled from two independent experiments. For MA subsp. *abscessus*, statistical time effect was evaluated with Mann-Whitney Test. Statistical comparison between MA subsp. *abscessus* and control mice at Day 7 was calculated with Mann-Whitney Test. *p<0.05, **p<0.01, ***p<0.001 ****p<0.0001.

Longitudinal monitoring by MRI was introduced to evaluate the progression of the pulmonary lesions within single mice at the day of infection and at time points 7, 45, 90 days. Lesions induced by MA subsp. *abscessus* were clearly visible at time points 45 and 90 days. In particular, infection foci appeared as a round well-defined area of consolidation with a preferential distribution in the left lung (**Figure 4**).

**Figure 4.**
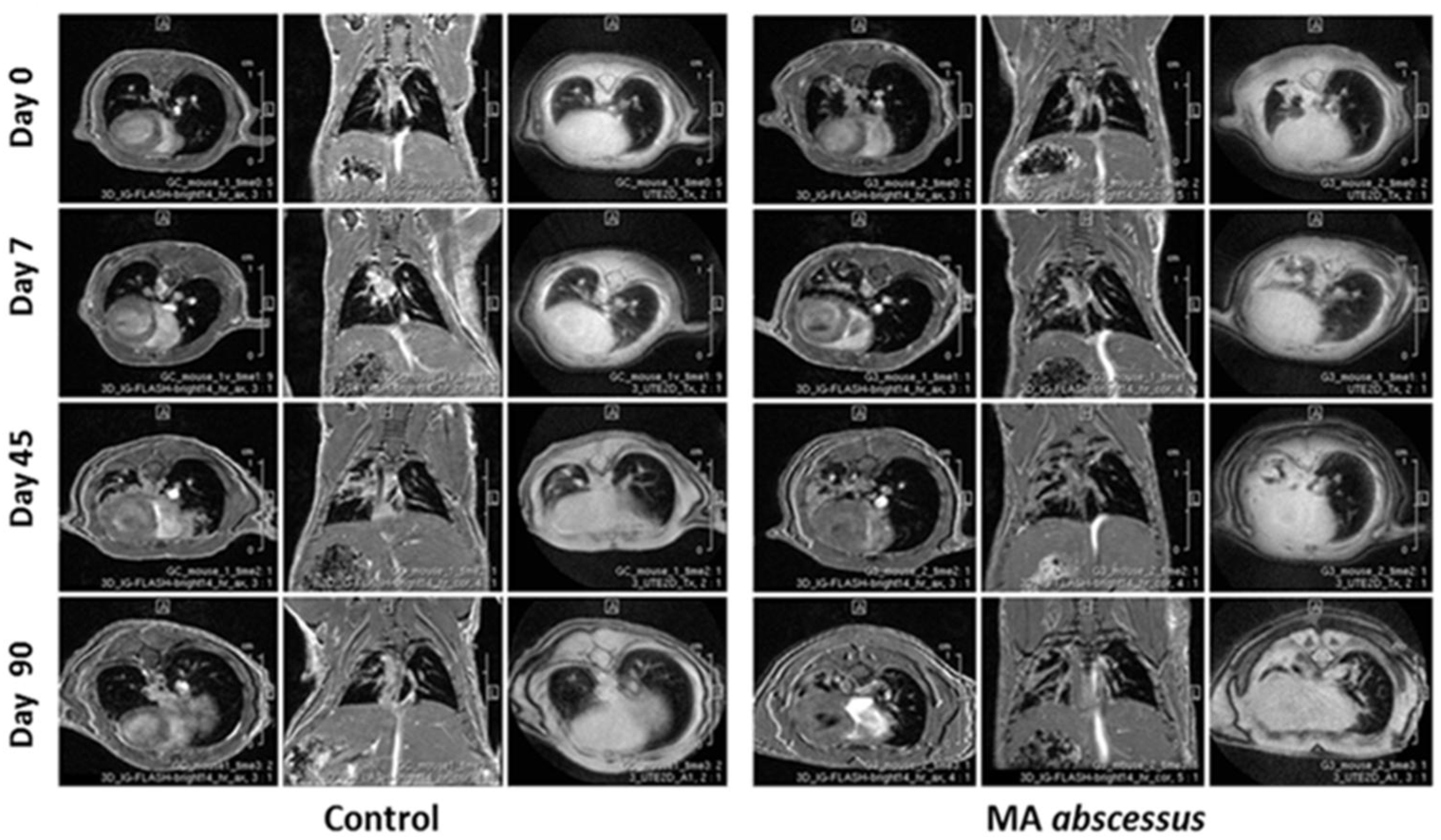
MRI analysis during the course of chronic MA subsp *abscessus* infection. Examples of representative axial and coronal 3D-FLASH and axial (2D-UTE) MR images of mice lungs obtained after inoculation with sterile agar beads (control mice) and with MA subsp. *abscessus*.

## 3. Discussion

The emerging problem of MA infections in the last 20 years has focused attention on the understanding of the pathogenesis of MA disease, which remains inadequately characterized [18]. Our aim was to develop an immunocompetent mouse model overcoming MA clearance and miming human MA airway disease (e.g. CF, COPD, bronchiectasis or asthma). We decided to intratracheally infect C57BL/6NCrl mice, known as resistant to mycobacterial infection, with MA entrapped in small agar beads. This method was refined by Bragonzi *et al.* and has been successfully used to study *P. aeruginosa, B. cepacia* and *S. aureus* infection and co-infections [15-17]. So far, C57BL/6NCrl mice were characterized by efficient clearance of MA from the lungs when inoculated via aerosol at a low dose; in contrast, airways clearance was much less effective following inoculation of high intravenous doses, ranging from 10^6^ to 10^8^ CFU [19, 20]. Embedding MA within agar beads and using intratracheal injection retained physically the bacteria in the airways and provided microanerobic/anaerobic conditions that allow bacteria to grow in microcolonies [21]. This procedure allowed us to create a stable infection in immunocompetent mice with all the three MA subspecies, and to reproduce a chronic infection useful for studying pathogenesis and progression of MA lung disease until three months from the inoculum. Statistical analysis detected the optimal breakpoint to distinguish high lung chronicity rate from low lung chronicity rate. The breakpoint was, for *MA* subsp. *abscessus*, 65 days and, for *MA* subsp. *bolletii* and *MA* subsp. *massiliense*, 45 days; after such time points, mice progressively inclined to clear the infection. Very recently *Le Moigne* [22] have published that intratracheal infection with agar beads is associated with very high mortality, in our hands the mortality has been very low and the chronicity was successfully established.

In human hosts MA can persist silently for years, and even decades, and can induce organized granulomatous lesions with foamy cells and in some cases, even caseous necrosis, like *M. tuberculosis* [23]. Infection with intracellular pathogens like MA is effectively controlled by the cell-mediated immune response and several cytokines play important roles limiting the host inflammatory response. In particular, IFN-γ is critical in the immune response to MA infection: it activates infected macrophages in concert with TNF-α and initiates a major effector mechanism of cell-mediated immunity. Immunocompetent mice, with evidence of strong TH1 response [24], appear resistant since they are characterized by a transient lung infection with a rapid clearance of MA in the first weeks [8-10]. Several attempts, with the aim of slowing down the resolution of MA lung infection, have led to establishing mouse models with deficits in innate or acquired immunity (e.g. GM-CSF – /, GKO or SCID mice) [11-13]. Moreover, corticosteroids were recently administered to immunocompetent mice to increase their susceptibility to the aereosolized pulmonary MA infection [25]. In our murine model, despite an initial protective high release of TNF-α and IFN-γ, granulomatous lung lesions (characterized by histiocytes, aggregates of foamy cells, neutrophils and lymphocytes) were present in the parenchyma around bronchi for about two months after MA infection, mimicking the damage observed in humans [23]. Interestingly, the presence of foamy cells, reported previously only in immunocompromised mice (SCID mice and GM-CSF -/- mice) [12] highly supports the reliability of this model with immunocompetent mice. For the first time, we introduced the monitoring of the MA lung infection with the MRI technique in this sudy. MRI represents a non-invasive approach providing an excellent qualitative evaluation for the assessment of inflammatory soft-tissue diseases in both humans and small laboratory animals [26], as well as an effective tool for monitoring lung inflammatory processes [27], without the possible detrimental impact on the immune system related to the use of X-ray based imaging approaches like Computed Tomography. MRI was applied in longitudinal measurements to monitor changes in infectious lesions, within the same C57BL/6NCrl mice, at the day of infection and after 7, 45, and 90 days from infection. Despite the decrease of pro-inflammatory markers and the dissolutions of granulomatous lesions during the course of the infection, the murine lung parenchyma was still damaged at 90 days as observed following both histopathological and MRI analysis, indicating a correlation between the 2 techniques. MRI was also able to discriminate, likewise the histopathological analysis, the lung inflammatory foci, induced by empty beads in control mice presenting as microscopic areas of consolidation at day 45 and 90.

MA is probably the most pathogenic of the NTMs infecting CF patients due to its multidrug resistance, poor response to treatment, and association with decline in lung function [28-30]. In CF patients MA pulmonary infections are associated with a wide clinical spectrum of disease, ranging from asymptomatic, transient colonization to significant lung function decline; making problematic the decision to start a combination therapy often associated with significant toxicity [30-32]. Published data show that the lack of suitable models of chronic infection jeopardise the urgent need to study the pathogenesis and the host immune response of MA lung disease. We show, for the first time, a long-term chronic infection model for all the three MA subspecies in immunocompetent C57BL/6NCrl mice, with similar clinical manifestations of the disease among the three subspecies as observed in humans [33]. We investigated the MA infection for up to six months when very few mice were still infected (data not shown) and the lung parenchyma was characterized by a residual peribronchial cuffing of CD3, B220, and IBA-1-positive cells (data not shown). Furthermore, we introduced the MRI technique for the first time in order to monitor the MA subspecies chronic lung infection over time by correlating imaging results with microbiological, immunological, and histological analyses. Our MA murine model could be adapted to study strain-related virulence in CF and non CF populations; representing a new suitable preclinical model for evaluating new antibiotics and innovative regimens.

## 4. Materials and Methods

### 4.1 Ethics statement

Animal studies were conducted according to protocols and adhering strictly to the Italian Ministry of Health guidelines for the use and care of experimental animals (IACUC N°816) and approved by the San Raffaele Scientific Institute Institutional Animal Care and Use Committee (IACUC).

### 4.2 Bacterial strains

MA subsp. *abscessus* (ATCC 19977), MA subsp. *bolletii* (ATCC 8156) and MA subsp. *massiliense* (ATCC 48898) were used. All the strains were conserved frozen at −80°C and were thawed and grown on Middlebrook 7H10 plates for three days at 37°C.

### 4.3 Mouse strains

Immunocompetent C57BL/6NCrl male mice (8 to 10 weeks of age) from Charles River were tested in the experiments. All mice were maintained in specific pathogen-free conditions at San Raffaele Scientific Institute, Milan, Italy.

### 4.4 Mouse model of chronic MA lung infection

Some colonies from 7H10 plates were grown for 2 days (to reach the exponential phase) in 20 ml of Middlebrook 7H9 broth. Then bacteria were embedded in agar bead preparation with modifications to the standard protocol [34]. The amount of white heavy mineral oil was reduced (50 ml) while the amount of Trypticase Soy Agar (TSA) was increased (25 ml). A lot of accurate washing steps were needed to remove traces of the mineral oil. For the inoculum, mice were anesthetized, the trachea was exposed and intubated, and 50 µL of beads suspension (1×10^5^ CFU) were injected before closing the incision with suture clips. Control mice were intratracheally inoculated with the same volume of empty beads suspension. After infection mice were daily monitored for body weight, appetite and hair coat; at fixed time points from infection (7, 14, 28, 45, 65 90 and 180 days). On average six mice were euthanized by CO_2_ asphyxiation.

Lung and spleen were collected and, once homogenized, were processed for microbiological analysis. Total CFU were the result of the addition of the CFU in lung homogenate and bronchoalveolar lavage fluid (BALF). The lung supernatants were stored at −80°C for cytokines and chemokines determination. In particular IFN-γ and TNF-α were evaluated by Mouse Custom ProcartaPlex 9-plex (Invitrogen, Thermo Fisher Scientific) and normalized at 2500 ug/ml of quantified proteins in lung supernatants.

### 4.5 Histological analysis and lung damage quantification

Formalin-fixed, paraffin-embedded sections of lungs at 7, 45, 90 and 180 days from MA infection were stained with H&E for histopathologic analysis. Immunohistochemical staining was performed with rabbit polyclonal anti-IBA-1 (Wako), monoclonal rat anti-human CD3 (Bio-Rad), rat anti-mouse B220/CD45R (clone RA3-6B2; Bio-Rad), after antigen retrieval. Immunoreactions were revealed by rabbit or rat on rodent HRP-polymer (Biocare Medical), using 3,3 diaminobenzidine (DAB) as chromogen (Biocare Medical) and slides were counterstained with haematoxylin. Ziehl-Neelsen staining was performed using automated Benchmark Special Stains instrument and dedicated staining kit (Roche). Photos of immunohistochemical staining and Ziehl-Neelsen staining were taken using AxioCam HRc (Zeiss) with the AxioVision System SE64 (Zeiss) while CF and Wt mice H&E were taken with Leica Biosystems Aperio and generated with Aperio ImageScope – Software.

Lung tissue lesions were quantified by Fiji ImageJ software [35] (version 1.52p) in 5 lobe images (Magnification 5X) of 5 mice for each time point (7, 45, 90 and 180 days from MA infection). In particular, the analysis was performed by Waikato Environment for Knowledge Analysis (WEKA), release v3.2.33 [36] (available at https://imagej.net/Trainable_Weka_Segmentation). The algorithm was trained to distinguish vessels/bronchi, lesions, healthy tissue, beads, blood, empty beads, artifacts/shadow (training set: at least 10 selections for each of the seven classes listed). Analysis was reported as the percentage of lesions to total lung lobe area.

### 4.6 MRI longitudinal monitoring

MRI studies were performed on a 7T preclinical scanner (Bruker, BioSpec 70/30 USR, Paravision 5.1), equipped with 450/675 mT/m gradients (slew-rate: 3400-4500T/m/s; rise-time 140µs) and a circular polarized mouse body volume coil with an inner diameter of 40 mm. MRI acquisitions were performed during gas anesthesia (Isofluorane, 3% for induction and 2% for maintenance 2L/min oxygen) and physiological monitoring. MRI scans were conducted with mice placed in a prone position inside the animal bed, scanning at fixed time points (7, 45, 90 and 180 days from MA infection). MRI protocol included: Three-Dimensional Intra-Gate Fast Low Angle Shot (3D IG FLASH) sequences acquired in the axial and coronal planes and a Two-Dimensional Ultra short Echo Time (2D UTE) sequences in axial plane. Sequence parameters were TR = 10 ms, TE = 2 ms, FOV = 25 × 24 mm, spatial resolution = 0,098 × 0,078 × 0,469 mm/pixel for 3D IG FLASH and TR = 30 ms, TE = 0, 478 ms, FOV 26 × 26 mm, spatial resolution = 0, 117 × 0, 117 mm/pixel, slice thickness 0,9 mm for 2D UTE.

### 4.7 Image analysis

Following acquisition, lung images were transferred to a dedicated workstation and analyzed with the use of MIPAV (Medical Image Processing, Analysis, and Visualization, National Institute of Health, Center for Information Technology). Images were analyzed by experienced radiologists. A region-of-interest (ROI) was drawn for each lung lesion on each slice on 3D-IG-FLASH axial images; the presence of the lesion was also confirmed by comparing coronal 3D-IG-FLASH and axial 2D UTE images, to avoid inclusion of potential artefacts. After segmentation of 3D-IG-FLASH axial images the whole volume of the affected lung parenchyma was automatically computed.

### 4.8 Statistics

A linear mixed-effects (LME)[37] model was employed to estimate the longitudinal trend of the body weight (log10 transformation) and evaluate the differences among groups (Control, MA subsp. *abscessus*, MA subsp. *bolletii* and MA subsp. *massiliense*). Different trends were allowed for Days ≤3 and Days >3. For Days >3, both linear and quadratic terms for time were included in the mixed models to account for the nonlinear trajectories of body weight over time. LME models were also employed to estimate the longitudinal trend of the bacterial load in the spleen and in the lung and the tissue damage. The *Ordered Quantile* normalization transformation of the outcome was considered in order to satisfy underlying model assumptions. Post-hoc analysis after LME was performed considering all the pairwise comparisons.

Logistic regression models were used to evaluate the lung chronicity rate during the course of the infection. The overall survival curves were estimated with the Kaplan-Meier method and were compared using the log-rank test. The Mann-Whitney test was performed to compare two independent groups, while in the presence of more than two independent groups the Kruskal-Wallis test followed by post-hoc analysis using Dunn’s test was used. Analyses were performed using R statistical software.

## Author Contributions

All authors have read and agreed to the published version of the manuscript.

## Abbreviations

NTMs: *Nontuberculous mycobacteria*
COPD: Chronic obstructive pulmonary disease
CF: Cystic fibrosis
MA: *Mycobaterium abscessus*
CFU: Colony forming units

