## Supplementary data for "A new model of chronic *Mycobacterium abscessus* lung infection in immunocompetent mice"

**Time course of MA subsp. *abscessus* chronic lung infection**

**Table S1**. Logistic regression model for chronicity (MA subsp. *abscessus,* Days 7-90).

| **Parameter** | **Estimate** | **SE** | **p-value** |
| --- | --- | --- | --- |
| Intercept | 5.8729 | 1.8978 | 0.0020 |
| Days | -0.0644 | 0.0244 | 0.0084 |

**Figure S1.** CFUs in total lung and spleen. Dots represent CFUs in individual mice (number of mice = 6-11). The data were pooled from two independent experiments. Linear mixed model followed by post hoc analysis was performed (**Table S2**). Significance was calculated for each time point comparing total CFU lung and total CFU spleen. The statistic comparison of total CFU lung among the different time point are in **Table S2**.

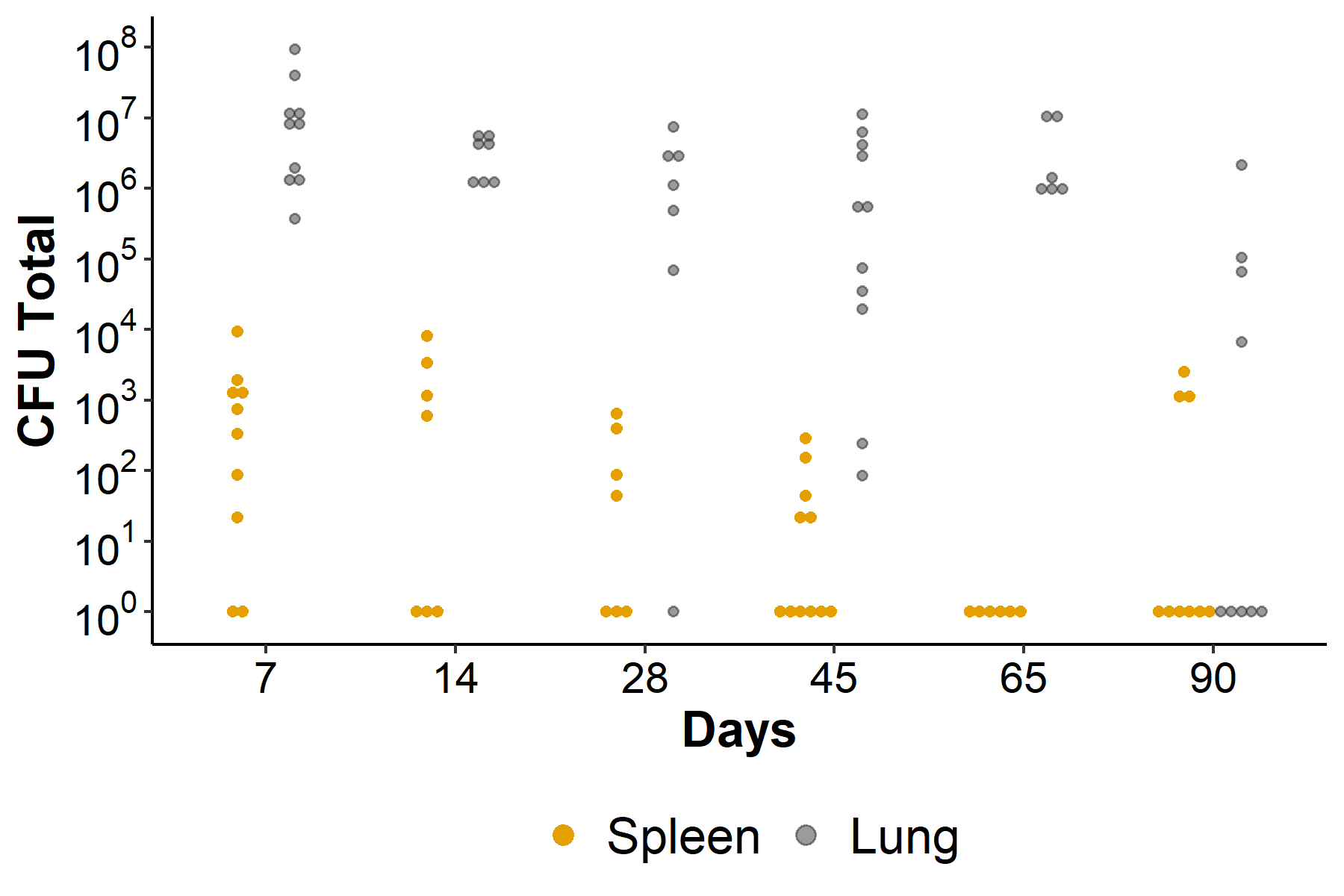

**Table S2**. Linear mixed model for MA subsp. *abscessus* total CFU lung and total CFU spleen (OrderNorm transformation). For testing differences between total CFU lung and total CFU spleen at each time point, a post-hoc analysis was performed.

| **Parameter** | **Estimate** | **SE** | **p-value** |
| --- | --- | --- | --- |
| Intercept | -0.1425 | 0.1542 | 0.3606 |
| Type (Ref=spleen) |  |  |  |
| Lung | 1.3843 | 0.1874 | **<0.0001** |
| Days (Ref=7) |  |  |  |
| Days14 | -0.1473 | 0.2403 | 0.543 |
| Days28 | -0.2908 | 0.2403 | 0.2327 |
| Days45 | -0.4318 | 0.2131 | 0.0488 |
| **Days65** | **-0.7447** | **0.2518** | **0.005** |
| Days90 | -0.4002 | 0.2241 | 0.081 |
| typeLung:Days14 | -0.122 | 0.2862 | 0.6721 |
| typeLung:Days28 | -0.1603 | 0.295 | 0.5898 |
| typeLung:Days45 | -0.1759 | 0.2548 | 0.4937 |
| typeLung:Days65 | 0.452 | 0.2995 | 0.1387 |
| **typeLung:Days90** | -1.113 | 0.2674 | **0.0004** |

| **Comparison** | **Days** | **Estimate** | **SE** | **p.value** |
| --- | --- | --- | --- | --- |
| Spleen – Lung | **7** | **-1.3843** | **0.1874** | **<0.0001** |
| Spleen – Lung | **14** | **-1.2623** | **0.2163** | **<0.0001** |
| Spleen – Lung | **28** | **-1.224** | **0.2278** | **<0.0001** |
| Spleen – Lung | **45** | **-1.2084** | **0.1725** | **<0.0001** |
| Spleen – Lung | **65** | **-1.8363** | **0.2336** | **<0.0001** |
| Spleen – Lung | 90 | -0.2713 | 0.1908 | 0.1623 |

**Figure S2. Time course of MA subsp. *bolletii* chronic lung infection*.*** C57BL/6NCrlBr mice were intratracheally infected with 1x10^5^ of MA *bolletii* reference strain (ATCC 8156) embedded in agar-beads. After 7, 14, 28, 65, 90 days from infection mice were sacrificed. (**A**) Rate of chronicity (number of mice = 5-9). The data were pooled from two independent experiments (**Table S3**). (**B**) Mice were considered infected with CFUs >0 in total lung . Dots represent CFUs in individual mice. The red line represents the median values. Kruskal-Wallis test with post-hoc multiple comparisons was performed (among the time points 7, 14, 28, 45 and 90) . *p<0.05, **p<0.01, ***p<0.001 ****p<0.0001,****p<0.0001. (**C**) CFUs in total lung and spleen. Linear mixed model followed by post hoc analysis was performed (**Table S4**). Significance was calculated for each time point comparing total CFU lung and total CFU spleen.

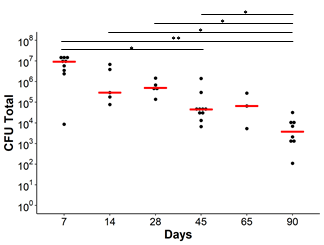
 A	B	C

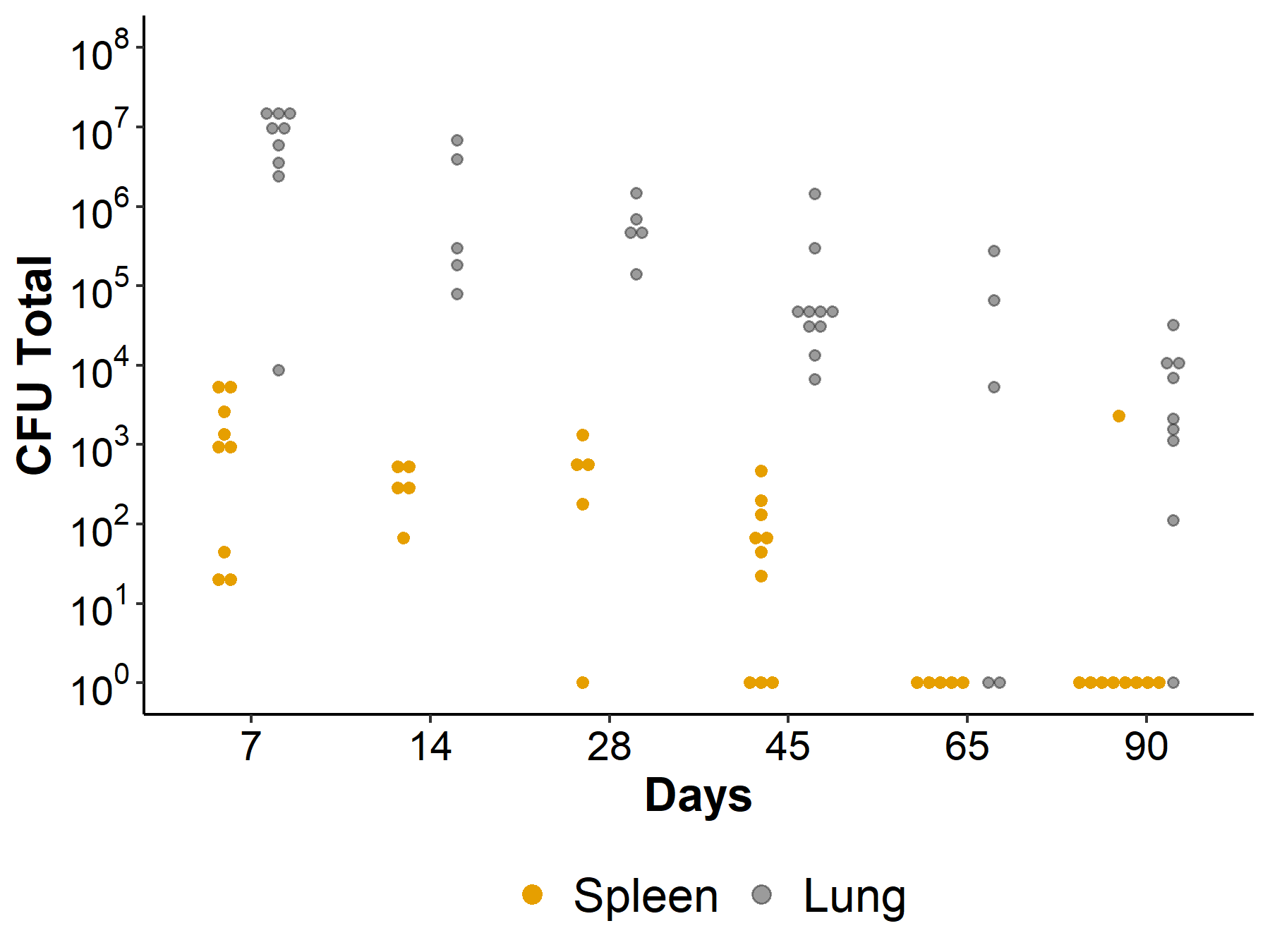

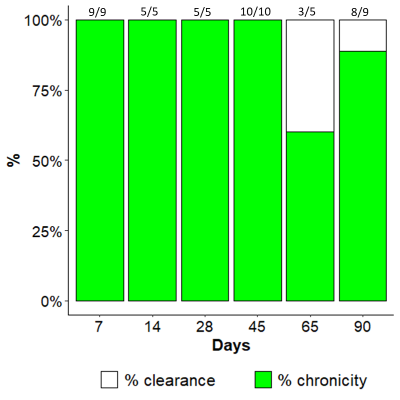

**Table S3**. Logistic regression model for chronicity (MA subsp. *bolletii*, Days 7-90).

| **Parameter** | **Estimate** | **SE** | **p-value** |
| --- | --- | --- | --- |
| Intercept | 4.8601 | 1.8843 | 0.0099 |
| Days | -0.0390 | 0.0249 | 0.1177 |

**Table S4**. Linear mixed model for MA *bolletii* total CFU lung and total CFU spleen (OrderNorm transformation). For testing differences between total CFU lung and total CFU spleen at each time point, a post-hoc analysis was performed.

| **Parameter** | **Estimate** | **SE** | **p-value** |
| --- | --- | --- | --- |
| Intercept | -0.0457 | 0.146 | 0.7558 |
| Type (Ref=spleen) |  |  |  |
| Lung | 1.6838 | 0.2065 | **<0.0001** |
| Days (Ref=7) |  |  |  |
| Days14 | -0.1275 | 0.2443 | 0.6049 |
| Days28 | -0.2252 | 0.2443 | 0.3626 |
| **Days45** | **-0.4915** | **0.2013** | **0.0195** |
| **Days65** | **-0.9995** | **0.2443** | **0.0002** |
| **Days90** | **-0.859** | **0.2065** | **0.0002** |
| typeLung:Days14 | -0.3837 | 0.3455 | 0.274 |
| typeLung:Days28 | -0.3406 | 0.3455 | 0.3306 |
| typeLung:Days45 | -0.4692 | 0.2846 | 0.1077 |
| typeLung:Days65 | -0.6647 | 0.3455 | 0.0621 |
| **typeLung:Days90** | **-0.6808** | **0.292** | **0.0253** |

| **Comparison** | **Days** | **estimate** | **SE** | **p.value** |
| --- | --- | --- | --- | --- |
| Spleen – Lung | 7 | -1.6838 | 0.2065 | **<0.0001** |
| Spleen – Lung | 14 | -1.3001 | 0.277 | **<0.0001** |
| Spleen – Lung | 28 | -1.3431 | 0.277 | **<0.0001** |
| Spleen – Lung | 45 | -1.2146 | 0.1959 | **<0.0001** |
| Spleen – Lung | 65 | -1.019 | 0.277 | **0.0007** |
| Spleen – Lung | 90 | -1.003 | 0.2065 | **<0.0001** |

**Figure S3. Time course of MA subsp. *massiliense* chronic lung infection*.*** C57BL/6NCrlBr mice were intratracheally infected with 1x10^5^ of MA *massiliense* reference strain (ATCC 48898) embedded in agar-beads. After 7, 14, 28, 65, 90 and 180 days from infection mice were sacrificed. (**A**) Rate of chronicity (number of mice = 5-15). The data were pooled from two independent experiments (**Table S5**). (**B**) Mice were considered infected with CFUs >0 in total lung. Dots represent CFUs in individual mice. The red line represents the median values. Kruskal-Wallis test with post-hoc multiple comparisons was performed (among the time points 7, 14, 28 and 45). *p<0.05, **p<0.01, ***p<0.001 ****p<0.0001,****p<0.0001. (**C**) CFUs in total lung and spleen. Linear mixed model followed by post hoc analysis was performed (**Table S6**). Significance was calculated for each time point comparing total CFU lung and total CFU spleen.

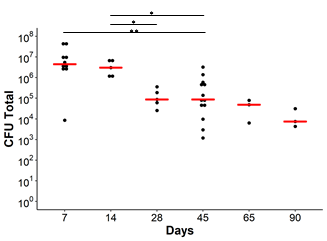
A	B	C

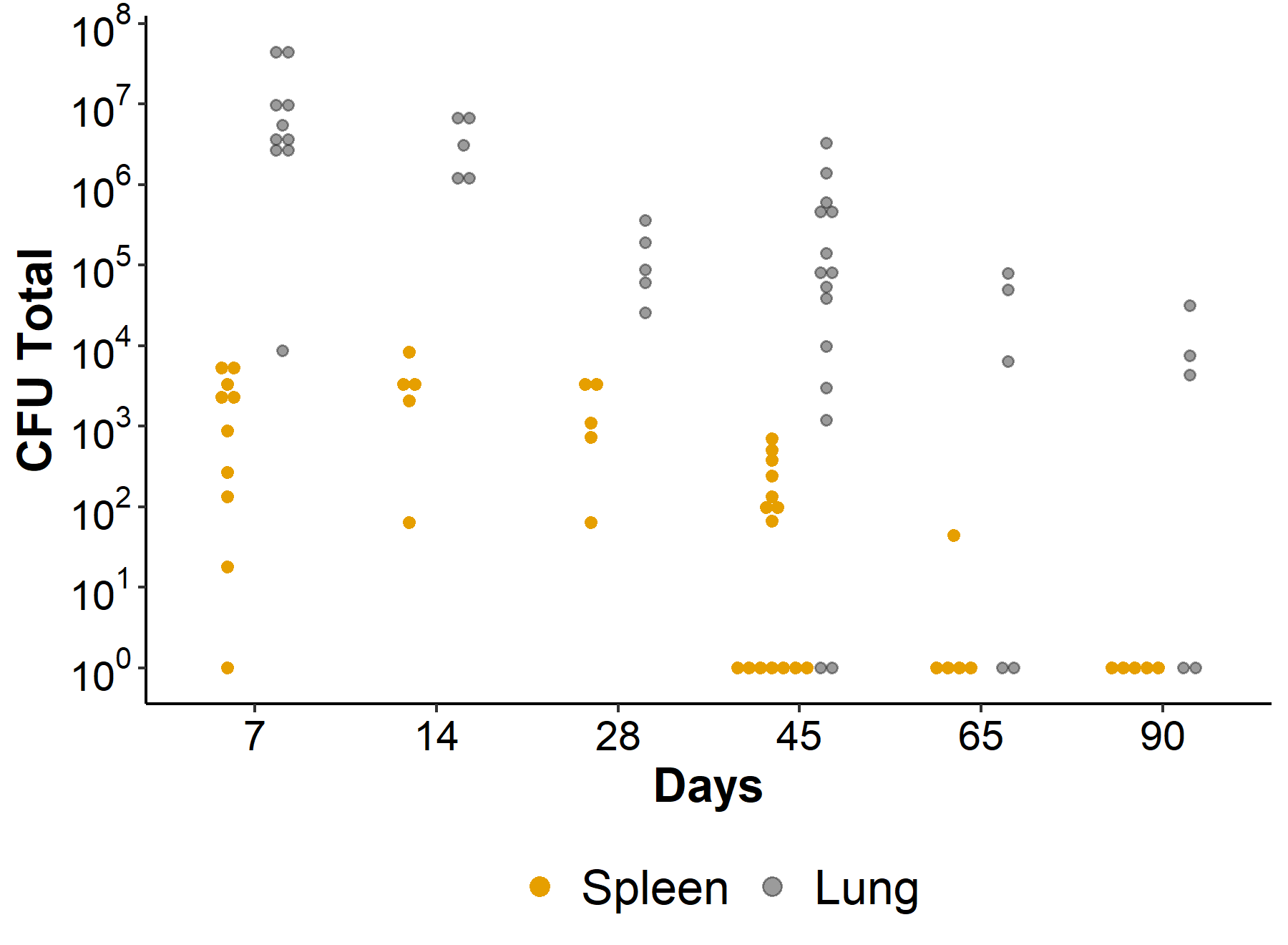

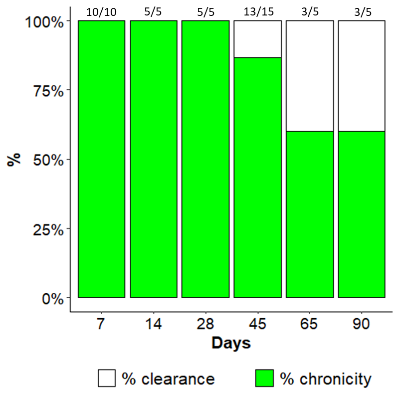

**Table S5**. Logistic regression model for chronicity (MA subsp. *massiliense*, Days 7-90).

|  | **Estimate** | **SE** | **p-value** |
| --- | --- | --- | --- |
| Intercept | 4.4041 | 1.3127 | 0.0008 |
| Days | -0.0504 | 0.0203 | 0.0133 |

**Table S6**. Linear mixed model for MA subsp. *massiliense* total CFU lung and total CFU spleen (OrderNorm transformation). For testing differences between total CFU lung and total CFU spleen at each time point, a post-hoc analysis was performed.

| **Parameter** | **Estimate** | **SE** | **p-value** |
| --- | --- | --- | --- |
| Intercept | -0.0873 | 0.1589 | 0.5859 |
| Type (Ref=spleen) |  |  |  |
| Lung | 1.6572 | 0.2089 | **<0.0001** |
| Days (Ref=7) |  |  |  |
| Days14 | 0.1792 | 0.2752 | 0.5189 |
| Days28 | 0.074 | 0.2752 | 0.7895 |
| **Days45** | **-0.4709** | **0.2052** | **0.0272** |
| **Days65** | **-0.7648** | **0.2752** | **0.0084** |
| **Days90** | **-0.8801** | **0.2752** | **0.0027** |
| typeLung:Days14 | -0.4138 | 0.3618 | 0.2597 |
| **typeLung:Days28** | **-0.9309** | **0.3618** | **0.014** |
| **typeLung:Days45** | **-0.6101** | **0.2697** | **0.0293** |
| **typeLung:Days65** | **-0.8664** | **0.3618** | **0.0215** |
| **typeLung:Days90** | **-0.8518** | **0.3618** | **0.0237** |

| **Comparison** | **Days** | **estimate** | **SE** | **p-value** |
| --- | --- | --- | --- | --- |
| Spleen – Lung | **7** | **-1.6572** | **0.2089** | **<0.0001** |
| Spleen – Lung | **14** | **-1.2434** | **0.2954** | **0.0004** |
| Spleen – Lung | **28** | **-0.7262** | **0.2954** | **0.0185** |
| Spleen – Lung | **45** | **-1.047** | **0.1705** | **<0.0001** |
| Spleen – Lung | **65** | **-0.7908** | **0.2954** | **0.0108** |
| Spleen – Lung | **90** | **-0.8053** | **0.2954** | **0.0095** |

**Figure S4**. Estimated model for the body weight (**Figure S3 A**, **Table S7**) and Kaplan–Meier estimates for overall survival (days) (**Figure S3 B, Table S8**) of mice.

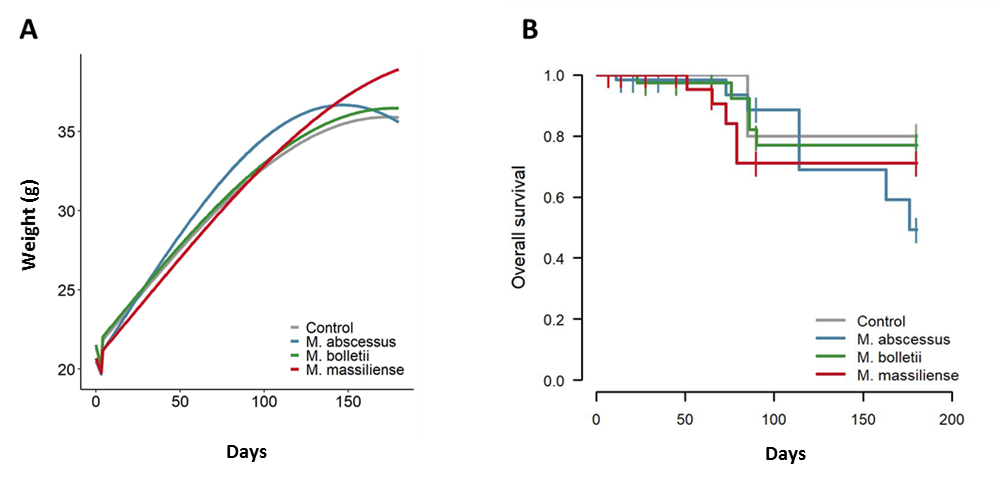

**Table S7**. Linear mixed-effects model for Body weight (log10 transformation). Different trends were allowed for time≤3 days (post infection) and time >3 days (post infection). For time >3 days, both linear and quadratic terms for time were included in the mixed models to account for the nonlinear trajectories of body weight over time.

|  | **Estimate** | **SE** | **p-value** |
| --- | --- | --- | --- |
| Intercept | 1.3288 | 0.0053 | <0.0001 |
| Type (Ref=Control) |  |  |  |
| MA. *abscessus* | -0.0173 | 0.0057 | 0.0028 |
| MA. *bolletii* | 0.0041 | 0.0059 | 0.4879 |
| MA. *massiliense* | -0.0131 | 0.0059 | 0.0262 |
| Days^2 | 0.0000 | 0.0000 | <0.0001 |
| Days*I(Days≤3) | -0.0067 | 0.0016 | <0.0001 |
| Days*I(Days>3) | 0.0026 | 0.0001 | <0.0001 |
| MA. *abscessus*:Days^2 | 0.0000 | 0.0000 | <0.0001 |
| MA. *bolletii*:Days^2 | 0.0000 | 0.0000 | 0.6717 |
| MA. *massiliense*:Days2 | 0.0000 | 0.0000 | 0.0041 |
| I(Days≤3):Days:MA. *abscessus* | 0.0008 | 0.0017 | 0.6142 |
| I(Days>3):Days:MA. *abscessus* | 0.0008 | 0.0001 | <0.0001 |
| I(Days≤3):Days:MA. *bolletii* | -0.0046 | 0.0018 | 0.0109 |
| I(Days>3):Days:MA. *bolletii* | 0.0000 | 0.0001 | 0.7792 |
| I(Days≤3):Days:MA. *massiliense* | 0.0001 | 0.0018 | 0.9447 |
| I(Days>3):Days:MA. *massiliense* | 0.0000 | 0.0001 | 0.9379 |

**Table S8.** Mortality rate statistics.

| **7 days post infection (p.i)** | **Total**  **Number** | **Survived**  **Number (%)** | **Dead**  **Number (%)** |
| --- | --- | --- | --- |
| Control | 12 | 12(100%) | 0(0%) |
| MA. *abscessus* | 74 | 74(100%) | 0(0%) |
| MA. *bolletii* | 54 | 54(100%) | 0(0%) |
| MA. *massiliense* | 56 | 56(100%) | 0(0%) |

| **7 days p.i - 14 days p.i.** | **Total**  **Number** | **Survived**  **Number (%)** | **Dead**  **Number (%)** |
| --- | --- | --- | --- |
| Control | 8 | 8(100%) | 0(0%) |
| MA. *abscessus* | 64 | 63(98.44%) | 1(1.56%) |
| MA. *bolletii* | 45 | 45(100%) | 0(0%) |
| MA. *massiliense* | 46 | 46(100%) | 0(0%) |

| **14 days p.i. 28 days p.i.** | **Total**  **Number** | **Survived**  **Number (%)** | **Dead**  **Number (%)** |
| --- | --- | --- | --- |
| Control | 8 | 8(100%) | 0(0%) |
| MA. *abscessus* | 49 | 49(100%) | 0(0%) |
| MA. *bolletii* | 40 | 39(97.5%) | 1(2.5%) |
| MA. *massiliense* | 41 | 41(100%) | 0(0%) |

| **28 days p.i. -45 days p.i.** | **Total**  **Number** | **Survived**  **Number (%)** | **Dead**  **Number (%)** |
| --- | --- | --- | --- |
| Control | 8 | 8(100%) | 0(0%) |
| MA. *abscessus* | 34 | 34(100%) | 0(0%) |
| MA. *bolletii* | 34 | 34(100%) | 0(0%) |
| MA. *massiliense* | 36 | 36(100%) | 0(0%) |

| **45 days p.i. 65 days p.i.** | **Total**  **Number** | **Survived**  **Number (%)** | **Dead**  **Number (%)** |
| --- | --- | --- | --- |
| Control | 8 | 8(100%) | 0(0%) |
| MA. *abscessus* | 23 | 23(100%) | 0(0%) |
| MA. *bolletii* | 24 | 24(100%) | 0(0%) |
| MA. *massiliense* | 21 | 19(90.48%) | 2(9.52%) |

| **65days p.i. -90 days p.i.** | **Total**  **Number** | **Survived**  **Number (%)** | **Dead**  **Number (%)** |
| --- | --- | --- | --- |
| Control | 5 | 4(80%) | 1(20%) |
| MA. *abscessus* | 20 | 18(90%) | 2(10%) |
| MA. *bolletii* | 19 | 15(78.95%) | 4(21.05%) |
| MA. *massiliense* | 14 | 11(78.57%) | 3(21.43%) |

| **90 p.i. - 180 p.i.** | **Total**  **Number** | **Survived**  **Number (%)** | **Dead**  **Number (%)** |
| --- | --- | --- | --- |
| Control | 4 | 4(100%) | 0(0%) |
| MA. *abscessus* | 9 | 5(55.56%) | 4(44.44%) |
| MA. *bolletii* | 6 | 6(100%) | 0(0%) |
| MA. *massiliense* | 6 | 6(100%) | 0(0%) |

**Figure S5.** H&E and Ziehl Neelsen (20X, AxioCam HRc Zeiss) stained sections of lungs after 45 days of MA *abscessus* infection; scale bar: 50 μm.

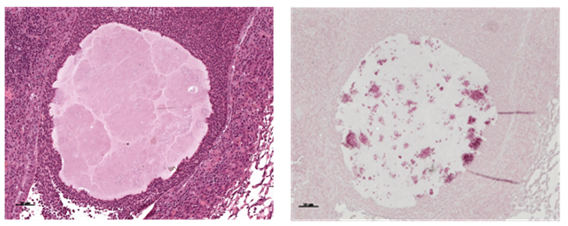

**Table S9.** Linear mixed model of tissue damage during MA subsp. *abscessus* chronic infection (square root transformation). Post-hoc analysis after LME was performed, comparing Control and MA *abscessus* at each time point.

| **Parameter** | | | **Estimate** | | **SE** | | **p-value** | | |
| --- | --- | --- | --- | --- | --- | --- | --- | --- | --- |
| Intercept | | | 0.1691 | | 0.0563 | | 0.0033 | | |
| Days (Ref=Days7) | | |  | |  | |  | | |
| Days45 | | | -0.0767 | | 0.0796 | | 0.3454 | | |
| Days90 | | | -0.0001 | | 0.0796 | | 0.9988 | | |
| Species (Ref=Control) | | |  | |  | |  | | |
| **MA abscessus** | | | **0.2318** | | **0.0796** | | **0.0077** | | |
| Days45:MA *abscessus* | | | -0.0370 | | 0.1126 | | 0.7456 | | |
| Days90:MA *abscessus* | | | -0.1155 | | 0.1126 | | 0.3152 | | |
| **Comparison** | | **Days** | | **estimate** | | **SE** | | | **p.value** |
| Control - MA. *abscessus* | | **7** | | **-0.232** | | **0.0796** | | | **0.0077** |
| Control - MA. *abscessus* | | **45** | | **-0.195** | | **0.0796** | | | **0.0221** |
| Control - MA. *abscessus* | | 90 | | -0.116 | | 0.0796 | | | 0.1572 |

**Figure S6. Lung lesions after 7, 45 and 90 days of chronic infection with MA subsp. *bolletii* (S6A*)* and MA subsp*. massiliense* (S6B)*.*** Dots represents the lobes analyzed by Image J program: 5 lobe images of 5 mice for each time point (25 dots for each time point). Total lung lesions were estimated in mice infected or not by MA subsp. *bolletii* and *massiliense*. Linear mixed model followed by post hoc analysis was performed (**Table S10 and S11**). Significance was calculated for each time point comparing MA subsp. *bolletii* and control and Ma subsp. *massiliense* and control *p<0.05, **p<0.01, ***p<0.001 ****p<0.0001.

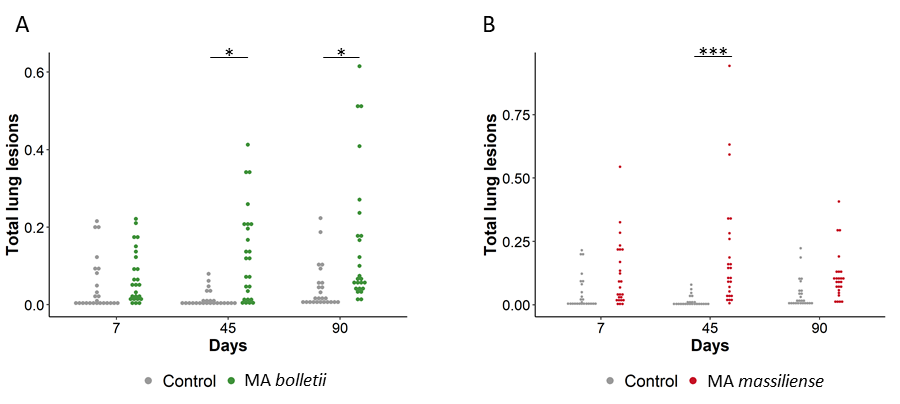

**Table S10.** Linear mixed model of tissue damage during MA subsp. *bolletii* chronic infection (sqrt transformation). Post-hoc analysis after LME was performed, comparing Control and MA *bolletii* at each time point.

| **Parameter** | | | **Estimate** | | **SE** | | **p-value** | | |
| --- | --- | --- | --- | --- | --- | --- | --- | --- | --- |
| Intercept | | | 0.1691 | | 0.0546 | | 0.0024 | | |
| Days (Ref=Days7) | | |  | |  | |  | | |
| Days45 | | | -0.0767 | | 0.0772 | | 0.3309 | | |
| Days90 | | | -0.0001 | | 0.0772 | | 0.9988 | | |
| Species (Ref=Control) | | |  | |  | |  | | |
| MA bolletii | | | 0.0687 | | 0.0772 | | 0.3829 | | |
| Days45:MA *bolletii* | | | 0.1475 | | 0.1092 | | 0.1895 | | |
| Days90:MA *bolletii* | | | 0.1116 | | 0.1092 | | 0.3171 | | |
| **Comparison** | | **Days** | | **estimate** | | **SE** | | | **p.value** |
| Control - MA. *bolletii* | | 7 | | -0.0687 | | 0.0772 | | | 0.3829 |
| Control - MA. *bolletii* | | **45** | | **-0.2162** | | **0.0772** | | | **0.01** |
| Control - MA. *bolletii* | | **90** | | **-0.1803** | | **0.0772** | | | **0.0283** |

**Table S11.** Linear mixed model of tissue damage during MA subsp. *massiliense* chronic infection (sqrt transformation). Post-hoc analysis after LME was performed, comparing Control and MA *massiliense* at each time point.

| **Parameter** | | | **Estimate** | | **SE** | | **p-value** | | |
| --- | --- | --- | --- | --- | --- | --- | --- | --- | --- |
| Intercept | | | 0.1691 | | 0.0550 | | 0.0026 | | |
| Days (Ref=Days7) | | |  | |  | |  | | |
| Days45 | | | -0.0767 | | 0.0778 | | 0.3344 | | |
| Days90 | | | -0.0001 | | 0.0778 | | 0.9988 | | |
| Species (Ref=Control) | | |  | |  | |  | | |
| MA massiliense | | | 0.1276 | | 0.0778 | | 0.1141 | | |
| Days45:MA *massiliense* | | | 0.1620 | | 0.1100 | | 0.1540 | | |
| Days90:MA *massiliense* | | | 0.0100 | | 0.1100 | | 0.9286 | | |
| **Comparison** | | **Days** | | **estimate** | | **SE** | | | **p.value** |
| Control - MA. *massiliense* | | 7 | | -0.128 | | 0.0778 | | | 0.1141 |
| Control - MA. *massiliense* | | **45** | | **-0.29** | | **0.0778** | | | **0.0011** |
| Control - MA. *massiliense* | | 90 | | -0.138 | | 0.0778 | | | 0.0898 |

**Figure S7.** Cytokines/chemokines recruitment during MA subsp. *bolletii* infection. (A) IFN-γ, (B) TNF-α measured by Mouse Milliplex, were quantified in total lung of mice. At 7 and 45 days dots represent cells in individual mice selected from the group of infected mice with MA subsp. *bolletii*. The data were pooled from two independent experiments. For MA subsp. *bolletii*, Statistical time effect was evaluated with Mann-Whitney Test. Statistical comparison between MA subsp. *Bolletii* and control mice at Day 7 was calculated with Mann-Whitney Test. *p<0.05, **p<0.01, ***p<0.001 ****p<0.0001.

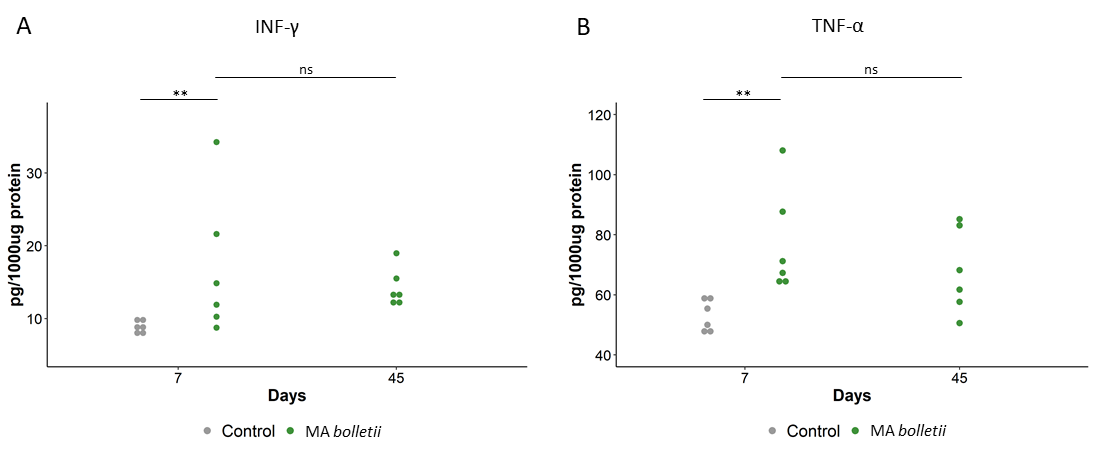

**Figure S8.** Cytokines/chemokines recruitment during MA subsp. *massiliense* infection. (A) IFN-γ, (B) TNF-α. measured by Mouse Milliplex, were quantified in total lung of mice. At 7 and 45 days dots represent cells in individual mice selected from the group of infected mice with MA subsp. *massiliense*. The data were pooled from two independent experiments. For MA subsp. *massiliense*, statistical time effect was evaluated Mann-Whitney Test. Statistical comparison between MA subsp. *massiliense* and control mice at Day 7 was calculated with Mann-Whitney Test. *p<0.05, **p<0.01, ***p<0.001 ****p<0.0001.

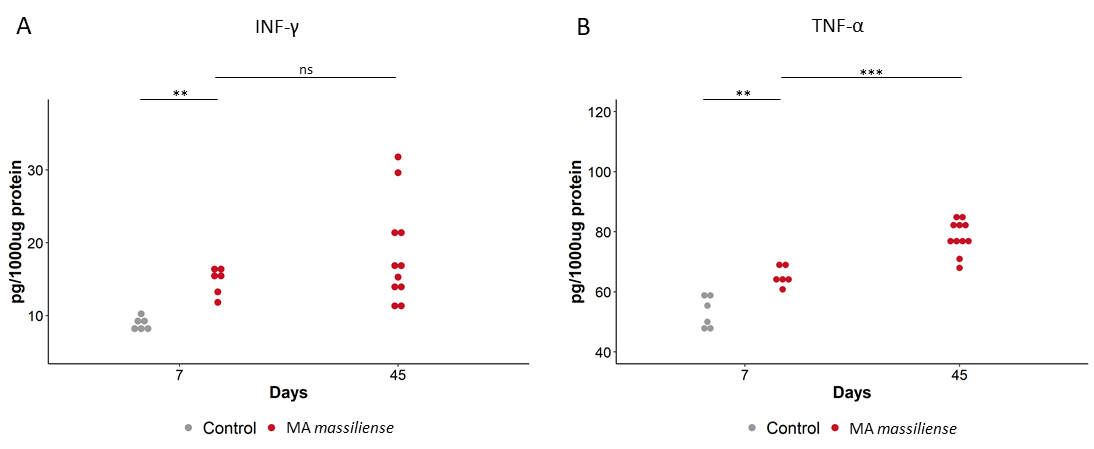
